# Effects of G2 checkpoint dynamics on the low-dose hyper-radiosensitivity

**DOI:** 10.1101/185371

**Authors:** Oluwole Olobatuyi, Gerda de Vries, Thomas Hillen

**Affiliations:** Department of Mathematical and Statistical Sciences, University of Alberta, Edmonton, Alberta, T6G 2G1, Canada.

**Keywords:** Linear Quadratic model, Induced Repair model, Early G2 checkpoint, Increased radioresistance, Hyper-radiosensitivity, Cell cycle, Cell cycle arrest, Mitotic catastrophe

## Abstract

We develop and analyze a system of differential equations to investigate the effects of G2 checkpoint dynamics on the low-dose hyper-radiosensitivity. In experimental studies, it has been found that certain cell lines are more sensitive to low-dose radiation than would be expected from the classical Linear Quadratic model (LQ model). In fact, it is frequently observed that cells incur more damage at a low dose (say 0.3 Gy) than at higher dose (say 1 Gy). This effect has been termed hyper-radiosensitivity (HRS). The HRS is followed by a period of relative radioresistance (per unit dose) of cell kill over the dose range of ~ 0.5 - 1 Gy. This latter phenomenon is termed increased radioresistance (IRR). These effects depend on the type of cells and on their phase in the cell cycle. Here we focus on the HRS phenomenon by fitting a model for the cell cycle that includes G2-checkpoint dynamics and radiation treatment to surviving fraction data for different cell lines including glioma cells, prostate cancer cells, as well as to cell populations that are enriched in certain phases of the cell cycle. The HRS effect is measured in the literature through 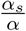, the ratio of slope *α*_*s*_, of the surviving fraction curve at zero dose to slope *α* of the corresponding LQ model. We derive an explicit formula for this ratio and we show that it corresponds very closely to experimental observations. Finally, we can identify the dependence of this ratio on the surviving fraction at 2 Gy. It was speculated in the literature that such a relation exists. Our theoretical analysis will help to more systematically identify the HRS in cell lines and opens doors to analyze its use in cancer treatment.

PACS and mathematical subject classification numbers as needed.

## 1. Introduction

The cell cycle is an ordered sequence of phases in the lifespan of a cell, which normally culminates in cell division. The cell cycle progression from one phase to the other is unidirectional. The four phases of the cell cycle are ordered as G1, S, G2, and M. The G1-phase is the first phase of a new daughter cell which lasts between 10-12h. This is the first growth phase where a cell increases its protein supply, increases the number of organelles like mitochondria and ribosomes, and increases in size. The S-phase, also referred to as the synthesis phase, starts when DNA replication commences. During this phase, which lasts about 5-7h, the amount of DNA in the cell effectively doubles. The G2-phase is the second growth phase lasting about 4h. This is the period of protein synthesis and rapid cell growth in preparation for cell division. The last phase is the M-phase, also referred to as the mitotic phase. This is the shortest cycle which last about 2h. During this stage, the cell divides into two daughter cells.

As a cell progresses through this cycle, the integrity of its genome is ensured and maintained by regulatory mechanisms called checkpoints [6]. These checkpoints, *seemingly* situated at the entrance to the next cycle phase, ensure that a cell does not progress into the next phase with unrepaired damage. Usually, when a cell sustains any form of damage in a particular phase, the checkpoint of that phase will stop the cell’s progression into the next phase in order to allow more time for damage repair. The process that stops cycle progression in order to give more time for repair is referred to as cell cycle arrest [24]. Moreover, as soon as the repair is complete, the cell is allowed to continue through the cell cycle.

The type of damage that is of particular interest in this paper is radiation-induced damage and the corresponding response of various cell cycle checkpoints. Sometimes, the radiation damage is not recognized by the checkpoint, and damaged cells proceed to the next phase un-repaired. Damaged cells in the G2 phase that evade the G2 checkpoint have been shown to experience cell death shortly afterward. This cell death, which prevents mutation as a result of un-repaired damage, is called mitotic catastrophe [3]. Experiments have shown that checkpoint evasion predominantly occurs when cells are exposed to low doses of radiation (mostly below 0.5 Gy) [11]. Mitotic catastrophe of G2-phase cells that evade the checkpoint has been shown to result in increased cell death at low doses of radiation. This phenomenon is called Hyper-Radio-Sensitivity (HRS). As the radiation dose increases and the corresponding damage is recognized by the checkpoint, there is cell cycle arrest of the damaged cells, which results in an apparent resistance to radiation. This low-dose resistance phenomenon is referred to as Increased Radio-Resistance (IRR). Experiments involving cells in different phases have also shown that the HRS/IRR phenomena are more exaggerated in the G2-phase cells as compared to cells in other phases [30, 19, 15, 13, 7].

The surviving fraction (SF) of cells is traditionally modelled by the Linear Quadratic (LQ) model given by

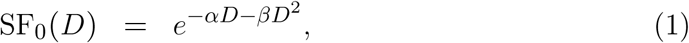

where *D* is the total radiation dose (measured in Gy), *α* is the rate at which single radiation tracks produce lethal lesions, and *β* is the rate at which binary misrepair of pairs of double strand breaks (DSB) from different radiation tracks lead to lethal lesions [1]. The LQ model is a monotonically decreasing function of dose; it cannot be used to describe the low-dose phenomena of HRS/IRR. In most cases, due to insufficient information about the low-dose cellular behaviors, the LQ model is usually extrapolated to low doses. However, experiments have shown that such extrapolation underestimates the low-dose radiation effects [13, 7].

In order to account for the HRS/IRR phenomena, Joiner et. al. [16] developed a modification of the LQ model, called the Induced Repair (IR) model, given by

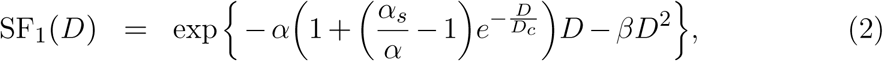

where *α*, *β*, and *D* are as defined in (1). *D*_*c*_ is the dose at which HRS transitions to the IRR phenomenon, and *α*_*s*_ is the initial slope of the surviving fraction curve (2) at *d* = 0 Gy. Figure 1 clearly explains these terminologies. Since the IR model was developed, the HRS/IRR phenomena have been widely quantified by the *α*_*s*_/ *α* index [10, 30]. It is easy to see that if 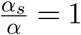, then the IR model reduces to the LQ model. Furthermore, any cell with 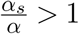 will exhibit the HRS/IRR phenomena.

**Figure 1:**
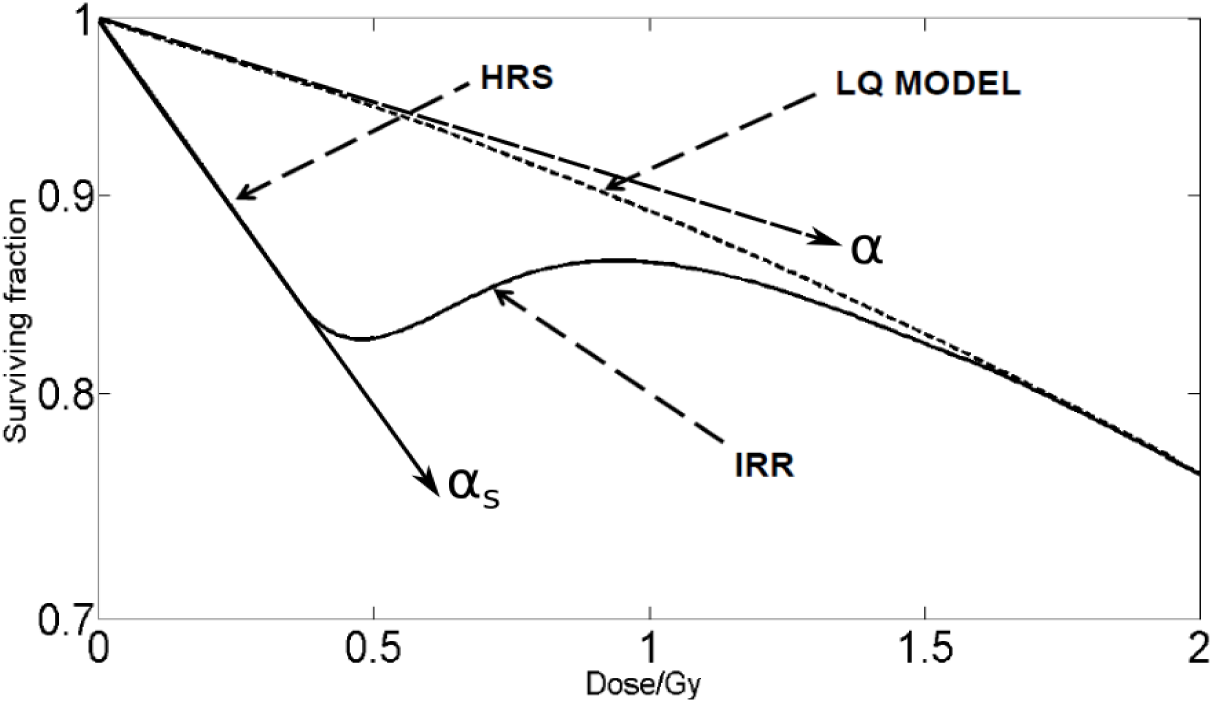
A surviving fraction curve illustrating the phenomena of HRS and IRR. The solid curve is the IR model while the dotted curve is the LQ model. *α*_*s*_ denotes the initial slope of the IR model while *α* denotes initial slope of the LQ model.

Experiments [12, 13] have been conducted at the molecular level in order to identify what is responsible for the HRS/IRR phenomena. It has been shown in [20, 15, 18] that the HRS/IRR phenomena consist of a sequence of cellular events such as checkpoint activity, mitotic catastrophe, cellular repair mechanism; and also the proportion of G2-phase cells at the time of radiation exposure. However, the relationship between the slope, *α*_*s*_, of the surviving fraction curve at *d* = 0 Gy and this sequence of events remains unclear. In particular, how can we interpret or model *α*_*s*_ in terms of this sequence of events?

In this paper, we focus on the HRS phenomenon by building an ordinary differential equations (ODE) model for the dynamics of cell cycle phases and damage repair, and how these are affected by radiation doses. We validate the model by fitting it to surviving fraction curves of ten asynchronous and three synchronous cell lines that exhibit the HRS phenomena. Then we derive the initial slope, *α*_*s*_, of the surviving fraction (SF) curve from this model for two cases motivated by the radiobiological experiments. In particular, we derive this initial slope of the SF curve when cell survival is computed immediately after radiation with (1) exposure time too short to accommodate cell progression and damage repair, and (2) exposure time long enough to accommodate damage repair but too short to accommodate cell cycle progression. Although in the experimental procedures leading to the measurement of the SF data, there is a post-radiation incubation period of 2 weeks during which survived cells can be recognized by their ability to form colonies (i.e., more than 50 daughter cells). This is because the current technology cannot detect survived cells until they are able to form colonies. However, since our deterministic model can compute the number of radiation-induced dead cells immediately after radiation exposure, we will assume that this model can be used as a proxy for measuring SF. Furthermore, experiments have shown that the repair of radiation damage usually occurs between 30-35mins after radiation [12]. Thus, since most SF data are measured from radiation exposure of at most 10 mins, we will use the derivation under the first assumption to compute the corresponding “*α*_*s*_” for our model. We find that the values of “*α*_*s*_” computed from our model agree with the data available in the literature. We also find the relationship between the initial slope *α*_*s*_, of the SF curve and some of the cellular events implicated in the HRS phenomena. Finally, we find an explicit relation of the 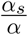 index to the surviving fraction at 2 Gy, which confirms speculations from the literature [14].

All through this paper, the term *synchronous* cell population will mean a culture of cells that is *rich* in a particular phase of the cell cycle. Thus, we will use this term interchangeably with the term cell culture *enriched* in a particular phase of the cell cycle. On the other hand, the term *asynchronous* cell population will refer to cell culture that is not enriched in a particular phase of the cell cycle.

The rest of this paper will be organized as follows: In Section 2, we formulate and develop the mathematical model from underlying cell cycle dynamics, and explain the relevant parameters. In Section 3, we fit the model to the surviving fraction data for 10 different asynchronous cell lines. We also fit the model to populations of cells enriched in various cell cycle phases. We estimate the model parameters and their 95% confidence intervals for these cell lines. In Section 4, we derive the analytical formula for the slope, *α*_*s*_, from the model under the two simplifying cases earlier mentioned. In Section 5, we numerically validate the derivations in the previous section. We find that the formula for the initial slope of SF curve derived in the previous section can replicate the values of *α*_*s*_ in the literature. In Section 6, we derive the relationship between 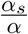 index and the radioresistance at 2 Gy. We find that cells with the same intrinsic radiosensitivity show an increasing relationship between 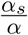 and the radioresistance at 2 Gy. This result contextualizes the suggestion of Lambin et. al [14]. We finish the paper with a discussion in Section 7.

## 2. Model formulation

Let *u* and *w* denote the population of cells in G2-and M/G1/S-phases, respectively. The model in this paper will be built from a simple model given by

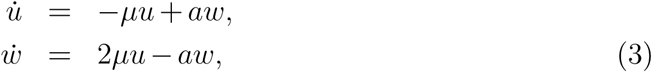

where *μ* is the rate of progression from G2 phase into mitosis, which results in two daughter cells entering the M/G1/S phase, and *a* is the rate of cell cycle progression from M/G1/S phase into G2 phase. We have restricted the model compartments to these two phase categories for simplicity. However, as we will see in Section 4, the derivation of the initial slope of the SF curve scales “nicely” with increase in the number of compartments.

We include radiation in the cell cycle dynamics (3), keeping in mind that the G2 checkpoint occurs in the damaged G2 phase. Thus, including the population of the damaged G2 cells, *v*, and the radiation terms along with their cellular interactions, we have,

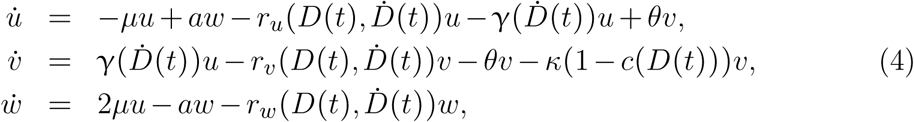

where *D*(*t*) denotes the total dose at time *t*, delivered at dose rate *Ḋ*(*t*), and is given by

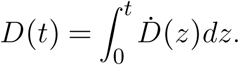

A schematic of the model, illustrating the relationship between the compartments, is given in Figure 2. When cells are exposed to radiation, a proportion of G2-phase cells is damaged at rate γ(*Ḋ*(*t*)). Damaged cells in the G2 phase undergo mitotic catastrophe at rate *κ*. The rate of mitotic catastrophe reduces when the G2 checkpoint, modeled by *c*(*D*), is activated. *c*(*D*) = 1 corresponds to a fully activated checkpoint and *c*(*D*) = 0 corresponds to the state when the checkpoint is not activated. The activation of this checkpoint arrests the damaged cells in the *v* compartment in order to give time for repair which occurs at a rate, *θ*.

**Figure 2:**
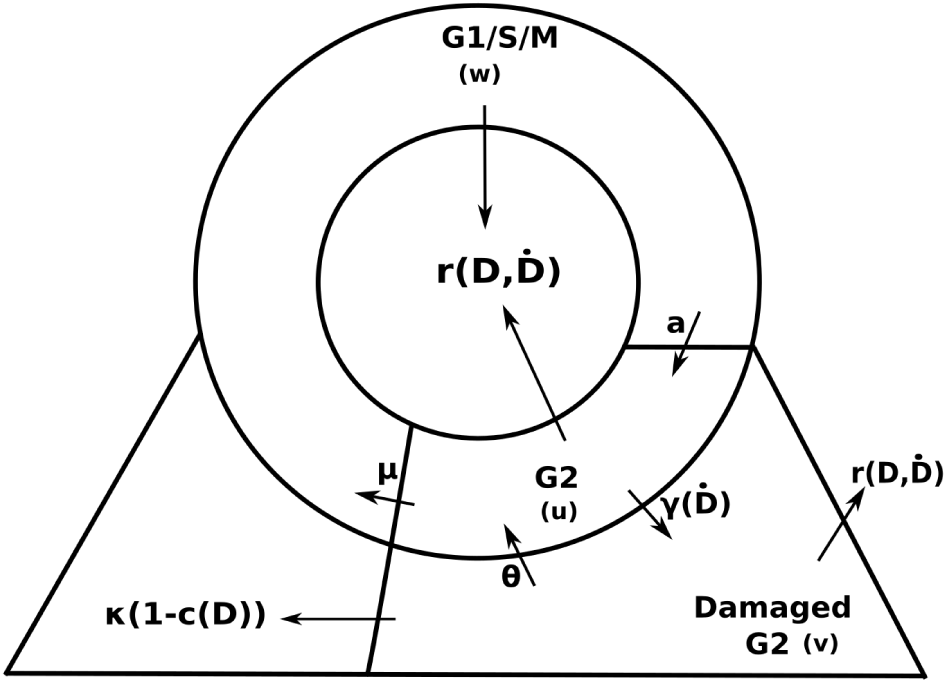
Schematics for the model in (4). The circle is partitioned into G2 and M/G1/S compartments. The arrows denote the rates of change between compartments. The damaged G2 compartment is an extension of the G2 compartments. The extension of the M/G1/S compartment denotes the damaged cells in mitosis that evade the G2 checkpoint.

We model the checkpoint dynamics by the function *c*(*D*), defined as

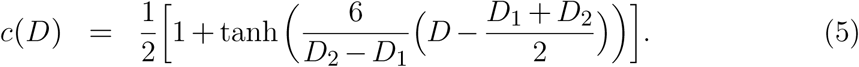

We denote *D*_1_ as the dose threshold below which the checkpoint fails to be activated (*c*(*D*) ≈ 0, ∀*D* < *D*_1_), and *D*_2_ as the dose threshold above which the checkpoint is fully activated (*c*(*D*) ≈ 1, ∀*D* ≥ *D*_2_) and damaged cells are arrested for repair. These thresholds are chosen to be in line with the experimental results in [12]. Figure 3 illustrates the profile of the checkpoint function *c*(*D*). This conforms to the hypothesis of Joiner et. al. in [11] which suggests that for HRS to occur at low doses, there must be a dose-sensing threshold below which damage is not sufficient to activate the repair mechanism. The half-saturation constant is given by 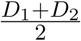 and the slope of the curve is given by 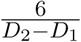. We are not particularly concerned whether the G2 checkpoint is activated as a result of more damaged cells or as a result of more damaged sites in a single cell. The dose-dependence of this checkpoint function, *c*(*D*), is sufficient to take care of these concerns.

**Figure 3:**
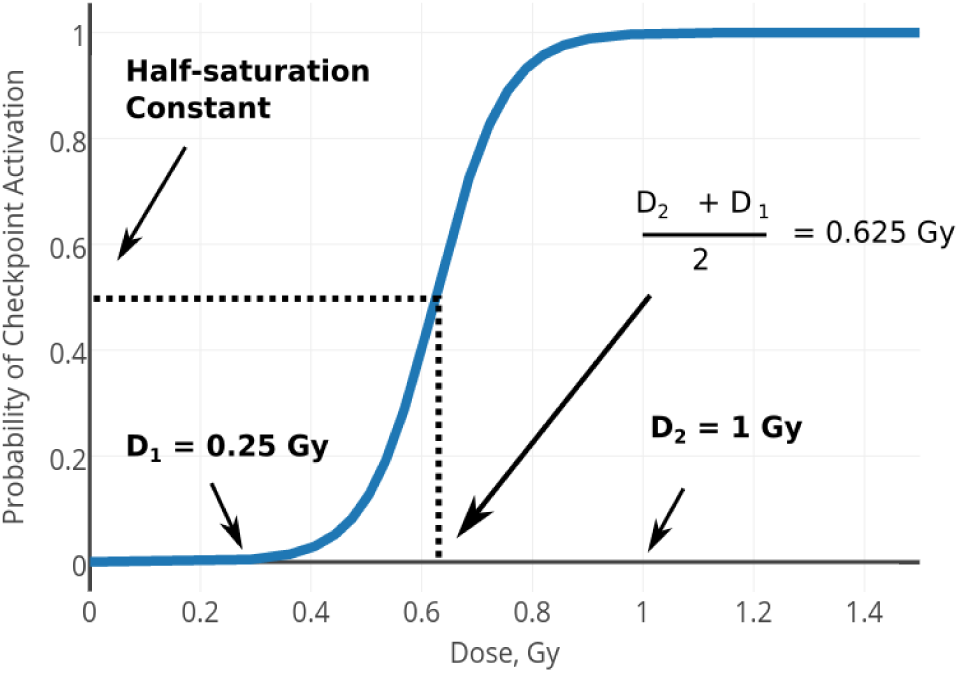
The profile of the G2 checkpoint functional. The lower threshold below which DNA is not recognized and the upper threshold above which the damaged cells are completely arrested for repair are given by *D*_1_ = 0.25 Gy and *D*_2_ = 1 Gy, respectively.

The two direct radiation effects on the cells are mainly the radiation-induced DNA damage and death, which occur at rates γ and *r*_*i*_, with *i* = {*u*, *v*, *w*}, respectively.

We model the radiation-induced damage in the G2 cells as proposed in [26], that is,

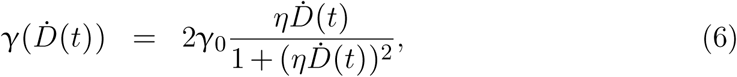

where γ_0_ is the maximum damage rate and 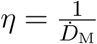 denotes the reciprocal of the dose rate *Ḋ*_*M*_ at which the radiation damage is maximal in a cell. We model the radiation-induced death rate by the radiation hazard function [8] given by

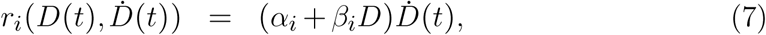

where *α*_*i*_ and *β*_*i*_ are as described in the LQ model for the different cycle phases, *i* = *u*, *v*, *w*.

We note here that the method we will use to derive the initial slope of the SF curve in Section 4 only works when the parameters and functions of the model (4) are independent of time. Therefore, we restrict the analysis to a time interval of constant radiation [0, *t*_0_] such that on this time interval the dose rate 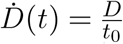 is constant. Since we are only interested in the first period of radiation exposure of length *t*_0_, we can drop the time dependence of (6)-(7) as follows:

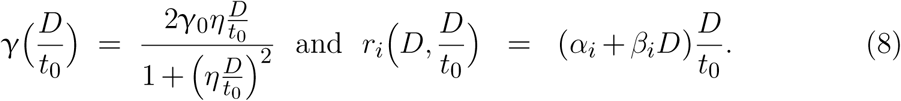

In the definition of *c*(*D*(*t*)), the checkpoint functional depends on the total dose given over total time, *t*_0_. Then,

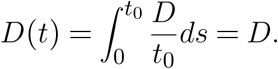

This implies that at time *t*_0_ we have

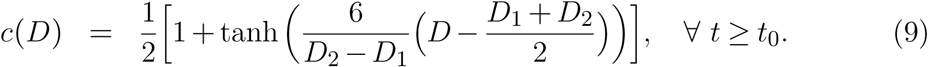

The data fitting in Section 3 and the derivation of the initial slope of the SF curve in Section 4 will be based on model (4) with time independent parameters and functions, namely

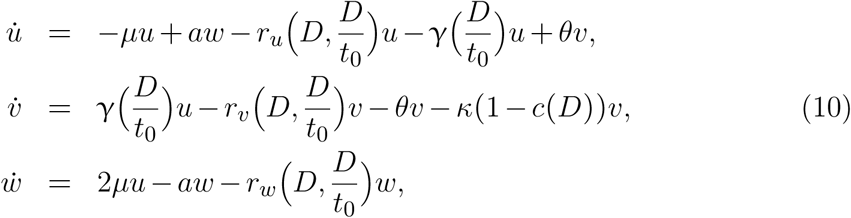

for 0 ≤ *t* ≤ *t*_0_. The terms 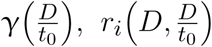, and *c*(*D*) are given by (8) and (9), respectively.

## 3. Data Fitting and Numerical Simulation

In this section, we fit a simplified version of model (10) to the surviving fraction data of different cell lines (both synchronous and asynchronous cells), and estimate the model parameters with their 95% confidence intervals. All of the cell lines in [12, 21, 16, 17, 17, 32, 32, 32, 30, 21] have been shown to exhibit the low-dose HRS phenomenon, although with varying degrees.

We will define the surviving fraction in the context of model (4). This is crucial because the formulas for the surviving fraction by both the LQ and the IR models do not account for the duration of the radiation exposure. The duration of the radiation exposure will be very significant in our analysis. Thus, we give the following definition:

#### Definition 1.

The surviving fraction *SF*(*D*, *t*) of cells at total dose *D* and time *t* from model (10) is given by

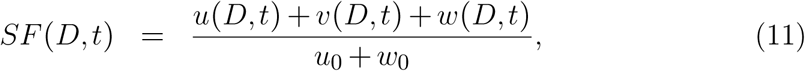

where *u*(*D*, *t*), *v*(*D*, *t*), and *w*(*D*, *t*) are the cell populations at time *t* and after radiation dose *D* in G2, damaged G2, and M/G1/S phases, respectively. We denote *u*_0_ = *u*(0,0) and *w*_0_ = *w*(0,0) as the initial populations of cells in G2 and M/G1/S phases, respectively, before radiation exposure started.

We fit the surviving fraction *SF*(*D*, *t*_0_) described in (11), to SF data over a radiation exposure time *t*_0_ chosen to be too short to accommodate both cell cycle progression and damage repair. This will be among the cases we will consider in the next section. As earlier noted, we will assume that the SF computed from model (10) is a good approximation to the experimental values of SF. Since the duration of radiation exposure is very short, we will also assume that the cell cycle progression rates *μ*, *a* and the DNA repair rate *θ* are zero in the simulation.

Thus, the simplification of model (10) we will fit to the asynchronous data, for 0 ≤ *t* ≤ *t*_0_, is given by

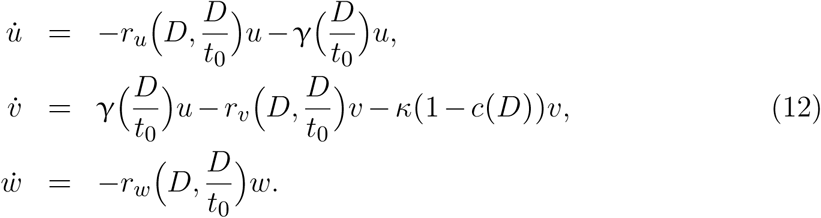

For the synchronous cell population, model (12) will be extended to accommodate all the phases of the cell cycle. Thus, for 0 ≤ *t* ≤ *t*_0_, we have

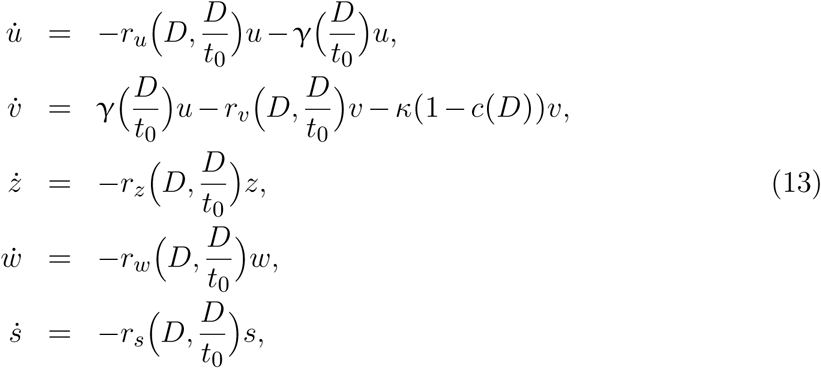

where *u*, *v*, *z*, *w*, *s* denote the population of cells in healthy G2, damaged G2, M-, G1-, and S-phases, respectively. We have also excluded the cell cycle rates, *μ*, *a*, and the DNA repair rate *θ* because the duration of radiation exposure *t*_0_ of interest is assumed to be too short to accommodate such dynamics.

In particular, we are interested in using the derived formula (26) in Section 4 to estimate the initial slope of the SF curve of these cell lines using the parameter values for γ_0_, *κ*, *η*, *D*_1_, *D*_2_, *α*, *β*, that will be estimated in this section. Notice that we have dropped the subscripts in the *α*’s and *β*’s in the data fitting. This is because in most of the literature, it is usually assumed that *α* ≔ *α*_*i*_ and *β* ≔ *β*_*i*_ for all *i* ∈ {*u*, *v*, *w*} in the asynchronous cell population. Similarly for the synchronous cell population, we will also assume that *α* ≔ *α*_*i*_ and *β* ≔ *β*_*i*_ for all *i* ∈ {*u*, *v*, *z*, *w*, *s*} and that *α* and *β* parameters will denote the radio-responsiveness of the enriched phase of the cell cycle. The ultimate goal is to compare these computed values of initial slope of the SF curve to the data in Figure 2 in [11].

The surviving fraction data of the various cell lines considered in this paper [12, 21, 16, 17, 17, 32, 32, 32, 30, 21] were measured from an experiment where cells were irradiated with single doses of X-rays between 0.05 and 6 Gy at dose rates ranging between 0.2-0.5 Gy/min. The surviving fraction of cells after exposure to a single dose was measured using a Cell Sorter. Most data points represent multiple measurements and are denoted as mean ± standard deviation. Cell survival was described in terms of their ability to form a colony (i.e., reproduce at least 50 offspring after radiation exposure) and cells which are unable to form a colony do not survive.

In this data fitting, we employ an implementation of the Goodman and Weare Affine invariant ensemble Markov Chain Monte Carlo (MCMC) sampler [9] to fit the model to the available surviving fraction datasets. The affine invariance property of this routine enables a much faster convergence even for badly scaled problems. This implementation takes, as an input, a log-likelihood function of the experimental data and a log-prior of each parameter. We assume an exponential distribution for the surviving fraction data. We also assume a uniform distribution for the prior of each parameter over the prescribed intervals of biologically relevant values.

In the following subsections, we will respectively fit models (12) and (13) to the asynchronous and synchronous cell data.

### 3.1. Asynchronous cell lines

In this subsection, we fit model (12) to the SF data of the following asynchronous cell lines: MR4[12], PC3[21], V79[16], V79ox[17], V79hyp[17], A549[32], HT29[32], U1[32], T98G[30], and RWPE1[21] cells. As mentioned earlier, these data describe the survival of cells after exposure to a single dose of 240kvp X-rays radiation. Doses between 0.05 and 6 Gy were delivered at dose rates ranging between 0.2-0.5Gy/min. Since there is no specific detail of which dose was delivered at a particular dose rate, we assume that every dose was delivered over a period of 10 mins. This assumption is fair because standard radiation technique delivers doses with appropriate dose rate over 10 mins. We assume that these asynchronous cells consist of 63% G1 phase cells, 19.35% S phase cells, and 17.65% G2 phase cells. This assumption is based on the mean of the proportion of cells in each phase that constitute asynchronous cell population in [19]. The values of the parameters and their respective 95% confidence intervals from the data fitting are given in Table 1.

**Table 1:** Estimated parameter values and their respective 95% confidence intervals from fitting model (12) to the SF data of the listed asynchronous cell lines.

| Cell line | $\gamma_0$ | $\kappa$ | $\eta$ | $d_1$ | $d_2$ | $\alpha$ | $\beta$ |
| --- | --- | --- | --- | --- | --- | --- | --- |
| MR4 | 0.255 ± 0.009 | 0.205 ± 0.009 | 50.75 ± 1.287 | 0.105 ± 0.009 | 0.575 ± 0.009 | 0.042 ± 0.002 | 0.160 ± 0.018 |
| PC3 | 0.138 ± 0.049 | 0.115 ± 0.044 | 29.93 ± 6.450 | 0.370 ± 0.065 | 0.720 ± 0.208 | 0.113 ± 0.066 | 0.050 ± 0.030 |
| V79 | 0.081 ± 0.0441 | 0.137 ± 0.073 | 23.87 ± 9.663 | 0.086 ± 0.038 | 0.379 ± 0.199 | 0.158 ± 0.016 | 0.030 ± 0.012 |
| V79ox | 0.128 ± 0.076 | 0.091 ± 0.059 | 17.74 ± 8.078 | 0.506 ± 0.188 | 1.057 ± 0.303 | 0.120 ± 0.054 | 0.044 ± 0.028 |
| V79hyp | 0.066 ± 0.036 | 0.092 ± 0.068 | 25.35 ± 9.516 | 0.503 ± 0.190 | 1.052 ± 0.303 | 0.086 ± 0.031 | 0.003 ± 0.002 |
| A549 | 0.066 ± 0.036 | 0.040 ± 0.030 | 90.40 ± 12.31 | 0.155 ± 0.038 | 0.261 ± 0.060 | 0.119 ± 0.035 | 0.045 ± 0.026 |
| HT29 | 0.092 ± 0.029 | 0.055 ± 0.029 | 25.22 ± 8.776 | 0.425 ± 0.086 | 0.848 ± 0.184 | 0.033 ± 0.016 | 0.055 ± 0.018 |
| U1 | 0.058 ± 0.019 | 0.024 ± 0.007 | 149.99 ± 8.00 | 0.035 ± 0.008 | 0.600 ± 0.032 | 0.013 ± 0.004 | 0.017 ± 0.004 |
| T98G | 0.201 ± 0.065 | 0.165 ± 0.073 | 47.59 ± 5.657 | 0.110 ± 0.022 | 0.792 ± 0.392 | 0.135 ± 0.033 | 0.018 ± 0.009 |
| RWPE1 | 0.300 ± 0.016 | 0.190 ± 0.016 | 55.65 ± 7.570 | 0.015 ± 0.008 | 0.385 ± 0.318 | 0.085 ± 0.023 | 0.090 ± 0.030 |

**Figure 4:**
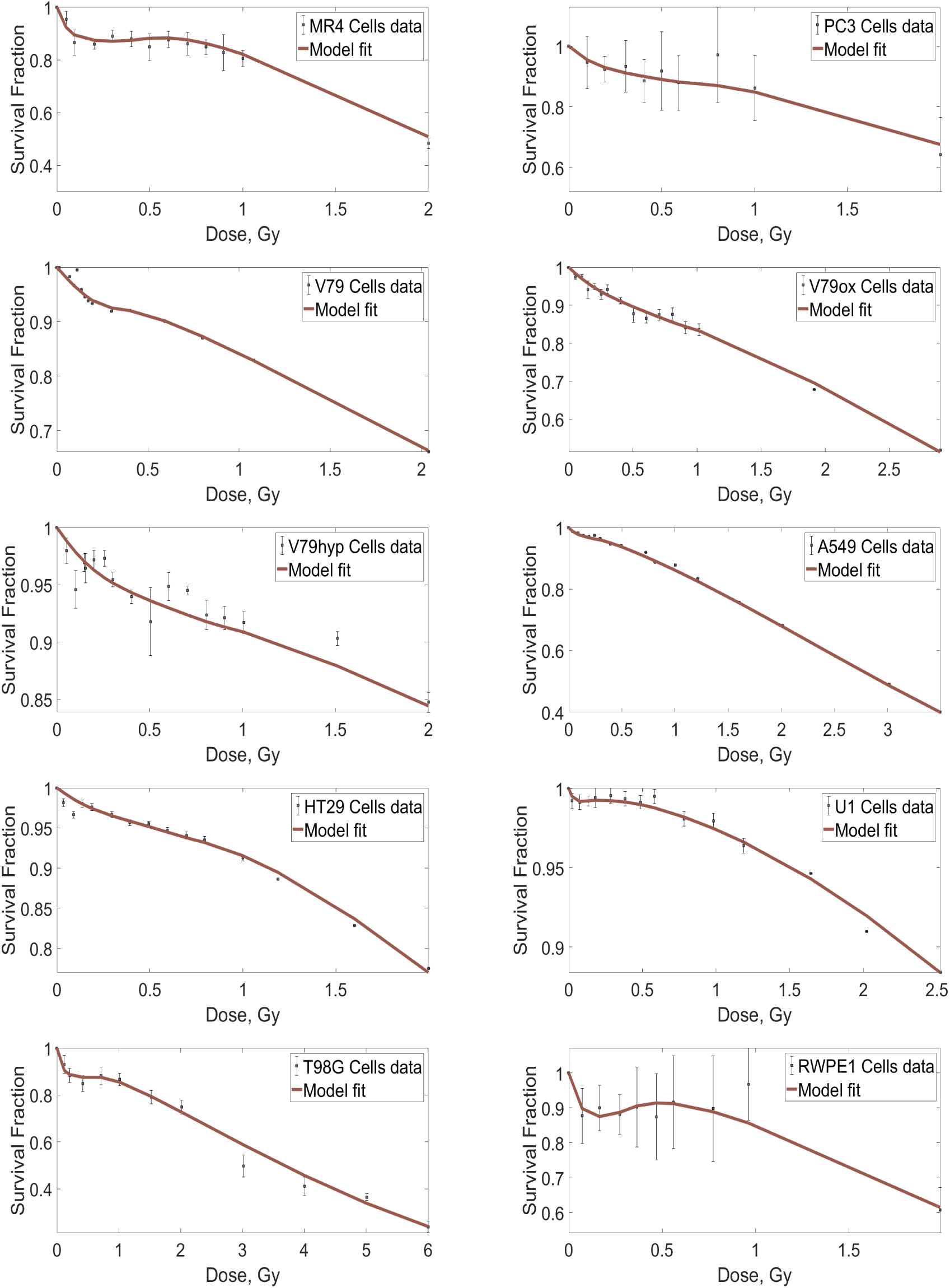
Data Fitting for the asynchronous cell population data. Model (12) is fitted to the surviving fraction data of ten different asynchronous cell populations. The data are shown with error bars and the best fit is shown in red.

### 3.2. Synchronous cell lines

In this subsection, we fit model (13) to the SF data of the V79, T98G, and U373 cell lines available for enriched G1, S and G2 phases [19, 30], respectively. The proportion of cells in different cell cycle phases at the time of radiation for the synchronous cell population are found in [19]. The estimated parameter values in this subsection are in Table 2 and the corresponding fits are in Figure 5.

**Table 2:** Estimated parameter values and their respective 95% confidence intervals from fitting model (13) to the SF data of the listed synchronous cell lines.

| Cell line | $\gamma_0$ | $\kappa$ | $\eta$ | $d_1$ | $d_2$ | $\alpha$ | $\beta$ |
| --- | --- | --- | --- | --- | --- | --- | --- |
| V79 G1 | 0.159 ± 0.079 | 0.103 ± 0.062 | 23.77 ± 9.743 | 0.400 ± 0.058 | 0.700 ± 0.231 | 0.250 ± 0.013 | 0.052 ± 0.023 |
| V79 S | 0.162 ± 0.082 | 0.105 ± 0.064 | 23.67 ± 9.437 | 0.400 ± 0.059 | 0.698 ± 0.220 | 0.085 ± 0.007 | 0.025 ± 0.007 |
| V79 G2 | 0.160 ± 0.069 | 0.103 ± 0.059 | 23.60 ± 7.349 | 0.399 ± 0.042 | 0.695 ± 0.172 | 0.303 ± 0.036 | 0.026 ± 0.016 |
| T98G G1 | 0.185 ± 0.035 | 0.118 ± 0.041 | 65.02 ± 11.80 | 0.200 ± 0.046 | 0.964 ± 0.321 | 0.130 ± 0.023 | 0.005 ± 0.002 |
| T98G S | 0.185 ± 0.011 | 0.115 ± 0.033 | 64.89 ± 10.90 | 0.201 ± 0.046 | 0.9 ± 0.153 | 0.130 ± 0.022 | 0.014 ± 0.001 |
| T98G G2 | 0.151 ± 0.062 | 0.110 ± 0.062 | 55.20 ± 7.062 | 0.150 ± 0.061 | 0.701 ± 0.130 | 0.21 ± 0.014 | 0.003 ± 0.001 |
| U373 G1 | 0.091 ± 0.048 | 0.070 ± 0.028 | 145.13 ± 9.712 | 0.009 ± 0.003 | 0.291 ± 0.061 | 0.020 ± 0.011 | 0.111 ± 0.039 |
| U373 S | 0.100 ± 0.042 | 0.071 ± 0.021 | 14.48 ± 1.069 | 0.010 ± 0.003 | 0.291 ± 0.045 | 0.100 ± 0.042 | 0.051 ± 0.023 |
| U373 G2 | 0.076 ± 0.033 | 0.065 ± 0.021 | 144.63 ± 11.53 | 0.009 ± 0.003 | 0.225 ± 0.038 | 0.160 ± 0.015 | 0.078 ± 0.004 |

**Figure 5:**
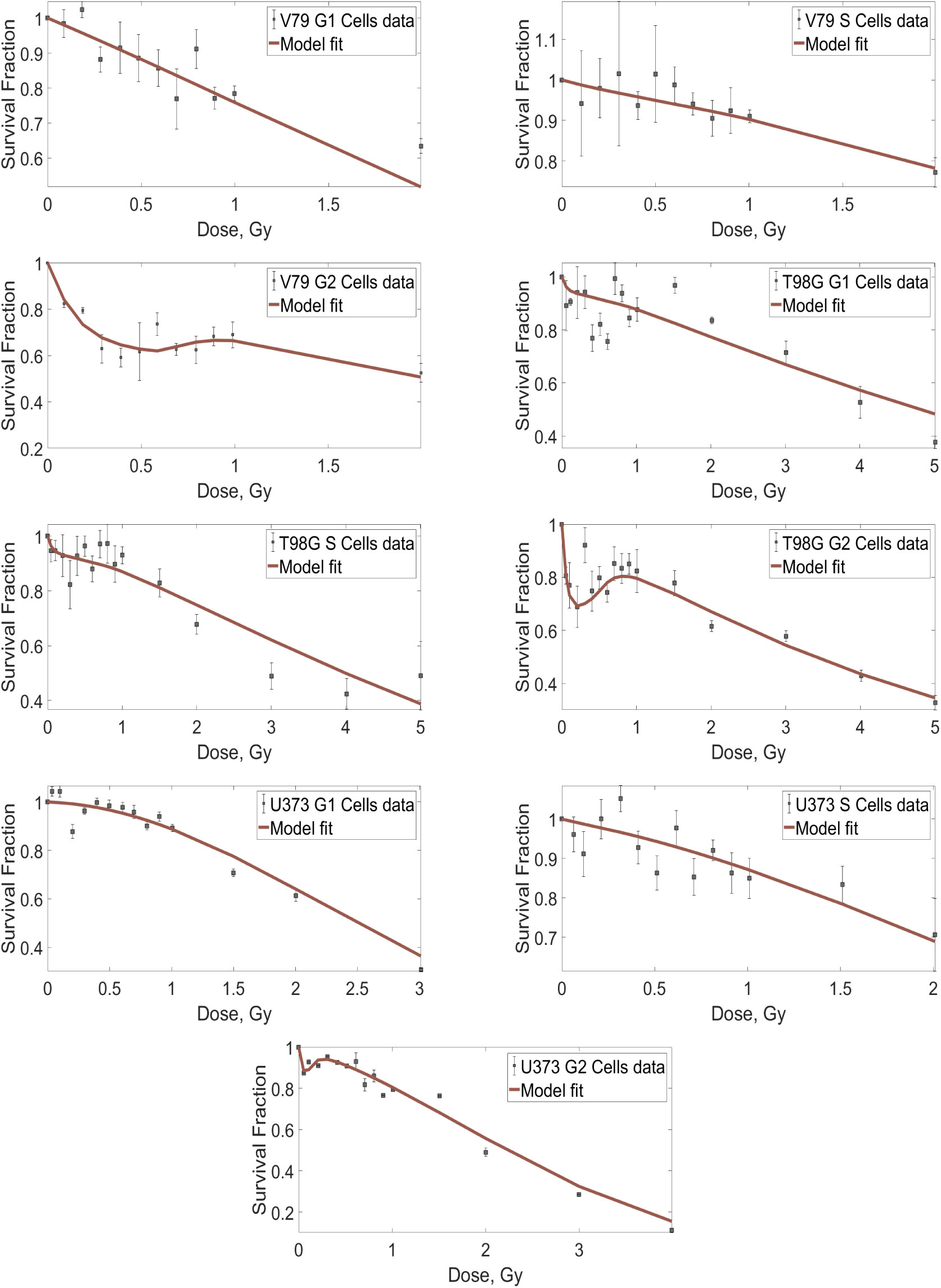
Data Fitting for the synchronous cell population data. Model (13) is fitted to the surviving fraction data of three different synchronous cell populations. The data are shown with error bars and the best fit is shown in red.

Notice that in Figure 5, some of other cell cycle phases aside from G2 also displays the phenomenon of HRS. For example, in the middle row of Fig. 5, both the enriched G1-and S-phase of the T98G cells exhibit a small measure of HRS. Our analysis shows that these enriched nonG2-phase cells that exhibit HRS contain a sizeable proportion of G2-phase cells, otherwise only enriched G2-phase cells has the capacity to exhibit these phenomena. For instance, the enriched G1-phase of both V79 and U373 cells was assumed to be in a ratio of 95.2% in G1, 4.7% in S, and 0.1% in G2 according to the data in [19]. Whereas the enriched G1-phase of T98G was assumed to be in a ratio of 85% in G1, 4.7% in S, and 10.3% in G2. These higher proportion of G2-phase cells in the enriched G1-phase of T98G explains the observed HRS which was not exhibited in both enriched G1-phase of both V79 and U373 cells. The same explanation goes for the S phase cells. This suggests that if we can synchronize the cells to obtain pure non-G2 phase cells, we might be better convinced that HRS phenomenon is exclusive to the G2 cells.

## 4. Derivation of the initial slope of the SF curve from the model

In the previous section, we have numerically shown that the model we formulated in this paper can describe the low-dose HRS phenomenon observed in both synchronous and asynchronous cells. Moreover, in this section, we will derive the formula for computing the initial slopes of these surviving fraction curves.

From the LQ and the IR models in (1) and (2), we have

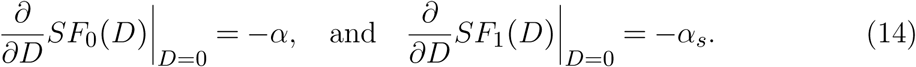

This implies that −*α* and −*α*_*s*_ are the initial slopes of the surviving fraction curve determined by both the LQ and the IR models, respectively. Hence, in order to derive the initial slope of the surviving fraction curve determined by the above model (10), it suffices to compute

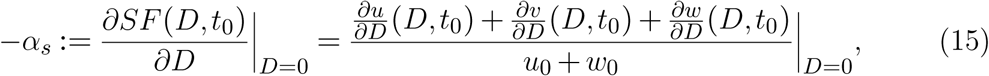

where 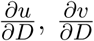, and 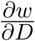 denote the partial derivatives of *u*, *v*, and *w*, with respect to *D. t*_0_ is the time at which the surviving fraction data is measured. Throughout this paper, the surviving fraction from the above model will be computed right after radiation exposure. Invariably, this implies that *t*_0_ is the radiation exposure time. In the following two subsections, we will derive the initial slope of the SF curve from model (12) for the two cases that *t*_0_ is

1. too short to accommodate both cell cycle progression and damage repair, and
2. too short to accommodate cell cycle progression but sufficient to accommodate damage repair.

The biological implications of these two cases will be discussed in Section 7.

### 4.1. Radiation exposure time too short for both repair of damage and cell progression

The model (10) can be written in vector form as

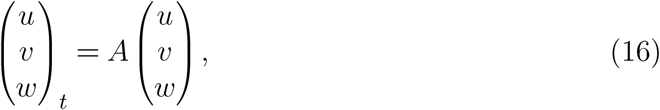

where

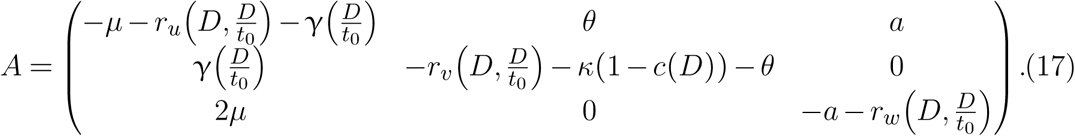

If we assume that the radiation time is too small to accommodate cycle progression and cell repair, matrix *A* then becomes

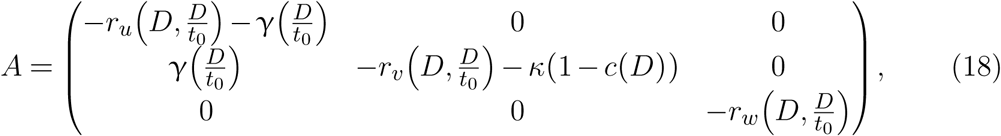

so that model (10) can be approximated by the reduced system in (12).

The system (12) can be solved explicitly with solution

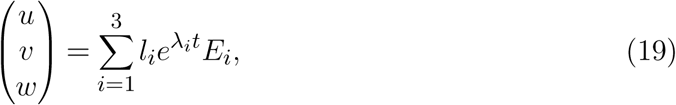

where λ_*i*_ is the eigenvalue of *A* that corresponds to eigenvector *E*_*i*_ for each *i* = {*u*, *v*, *w*}. To this end, let λ denote an eigenvalue of *A*. |*A* - λ*I*| = 0 implies

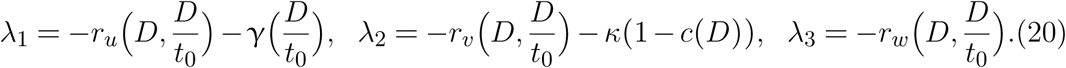

For each λ_*i*_, the corresponding eigenvector *E*_*i*_ is given by

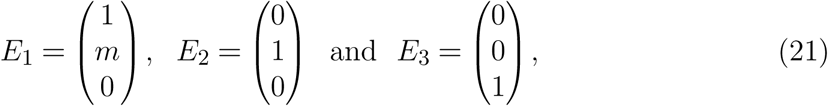

with

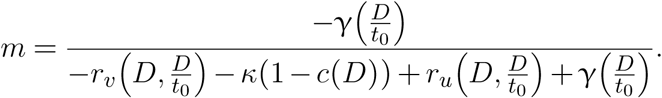

Thus, (19) with the corresponding eigenvalues and eigenvectors in (20) and (21) forms the general solution for the system (12). Now, using the initial conditions *u*(0) = *u*_0_, *v*(0) = 0, *w*(0) = *w*_0_, we have

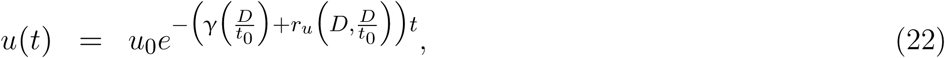

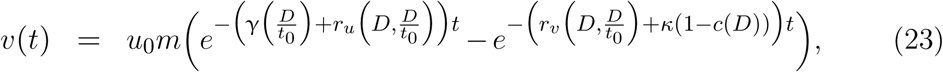

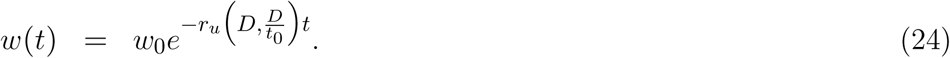

Since

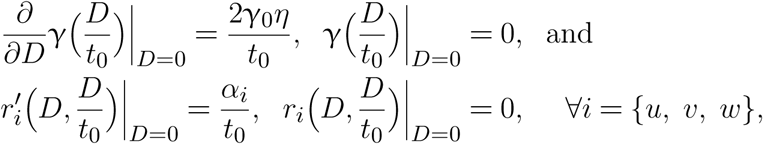

the partial derivatives of *u*, *v*, *w* with respect to *D*, evaluated at *D* = 0 and at time *t*_0_, are given by

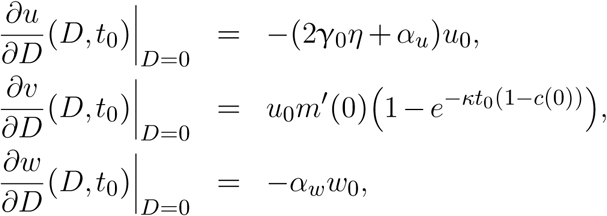

so that

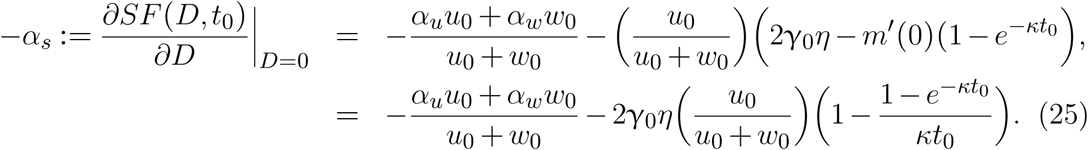

However, for asynchronous cell population where *α* ≔ *α*_*u*_ = *α*_*w*_, (25) simplifies to

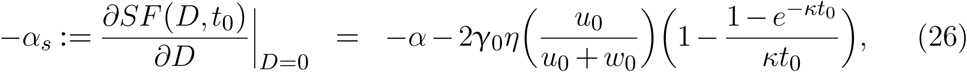

where the direct radiation effect is clearly described by *α*, which is the rate at which single radiation tracks produce lethal lesion in a general cell population. Moreover, for small radiation exposure time, *t*_0_, using Taylor expansion, we get a more compact form given by

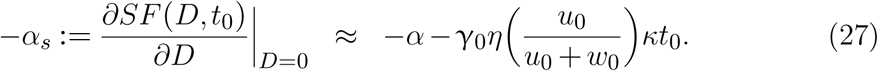

(27) reveals that the slope of the SF curve after a short duration of radiation exposure is controlled by two components, namely: the direct effect of radiation (captured by the first term on the RHS of (27)) and the indirect effect of radiation (captured by the second term on the RHS of (27)). The impact of this second term on *α*_*s*_ is controlled by the proportion of cells in G2-phase of the cell population. This is largely because the other parameters in the second term on RHS of (27) belong to the cells in the G2 phase of the cell cycle. This confirms the essence of the initial distribution of cells in various phases of the cell cycle at the time of radiation exposure on the phenomenon of low-dose HRS hypothesized in [11, 29, 19]. In fact, for synchronous cells enriched in G2-phase, the slope of the SF curve will be steeper because of the significant contributions from both the direct and the indirect radiation effects. On the other hand, for any synchronous cells enriched in non-G2-phase, the steepness of the slope is mainly controlled by the direct radiation effect *α* with very little contribution from the indirect radiation effect.

We also observe that the second term on the RHS of (27) is dominated by the parameter *κ*, denoting the rate of mitotic catastrophe. This implies that a major determinant of the degree of HRS exhibited in a cell is the rate of death of the G2-phase cells which evade the checkpoint activity.

### 4.2. Radiation exposure time too short for cell progression but sufficient enough for damage repair

In the previous subsection, we considered the case where the SF is computed right after radiation exposure whose duration is too short to accommodate cell cycle progression and DNA damage repair. Now we compute the surviving fraction immediately after a radiation exposure whose duration is long enough to accommodate repair but not sufficient to have any significant cycle progression. It is interesting to understand how the repair mechanism impacts the initial slope of the SF curve in this case. This problem is simply equivalent to solving (16) with *A* given by

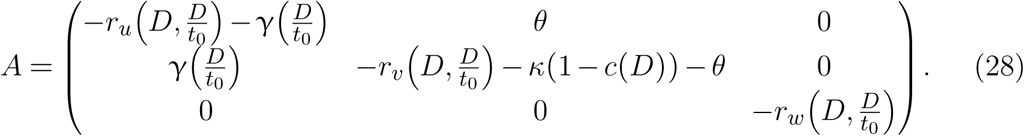

Notice that the difference between this matrix and the one in the previous subsection in (18) is the nonzero parameter *θ* in (28). As a check, this derivation should result in that of the last subsection when *θ* = 0.

Following the technique used in the last subsection, we can solve this system analytically to derive the following solution (where we have dropped the arguments for simplicity)

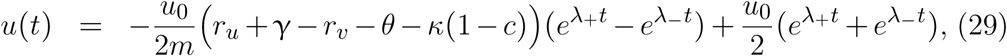

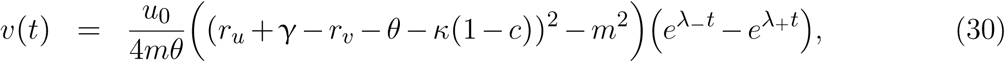

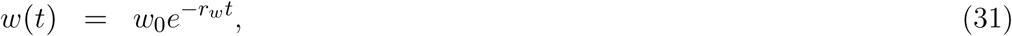

**Figure 6:**
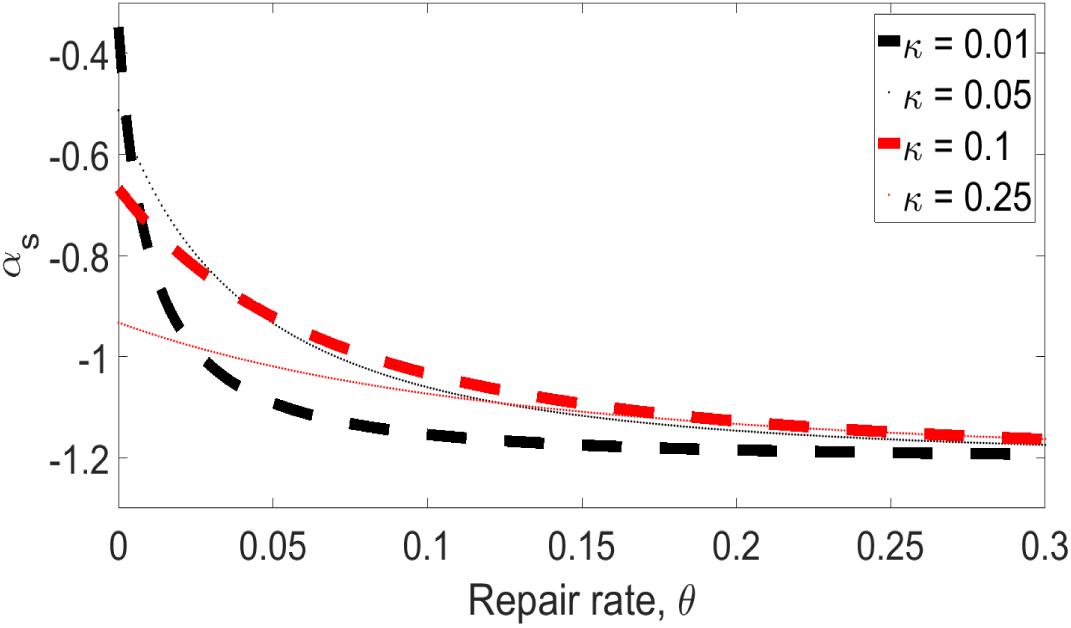
The profile of the initial slope of the SF curve as a function of repair rate *θ*, at different rates of mitotic catastrophe *κ*. The black dashed curve is for *κ* = 0.01, the black dotted curve is for *κ* = 0.05, the red dashed curve is for *κ* = 0.1, and the red dotted curve is for *κ* = 0.25.

where *u*_0_ = *u*(0,0) and *w*_0_ = *w*(0,0), with

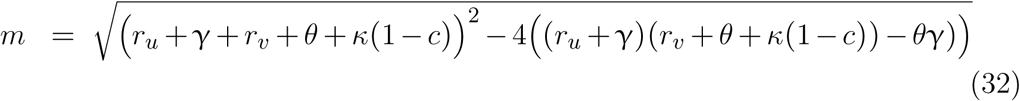

and

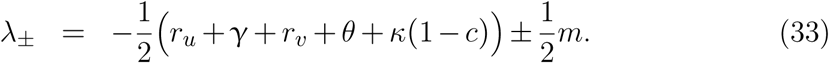

By computing 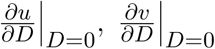, and 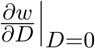, we can evaluate (15) to get

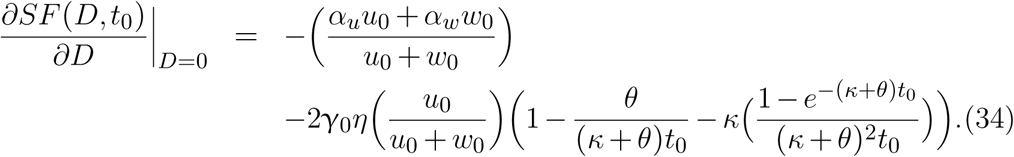

As before, assuming *α*_*u*_ = *α*_*w*_ = *α*, (34) reduces to

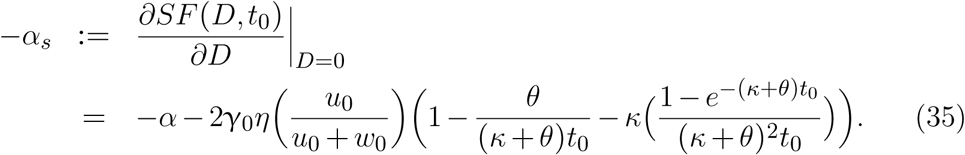

After Taylor expansion for small *t*_0_, we can get a more compact form given by

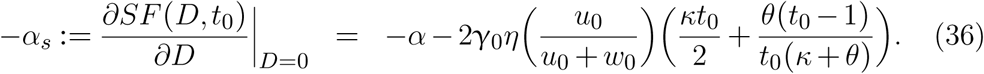

As noted earlier, we see that (27) is a special case of (35) when *θ* = 0. We can also understand the interaction between the rate of repairing DNA damage and the mitotic catastrophe experienced by damaged G2-phase cells that eventually culminates into the HRS phenomenon through (35). For small mitotic catastrophe rate *κ*, the slope *α*_*s*_ of the SF curve is more sensitive to changes in the DNA repair rate *θ* than for larger values of *κ*. Thus, the slope *α*_*s*_ of the SF curve depends on the relationship between the rates of mitotic catastrophe *κ* and DNA repair *θ*.

Furthermore, it is worth noting that (25) and (34) scale nicely with increase in the number of model compartments. For example, suppose model (10) has *n* compartments, where *u*_*i*_0__ and *u*_*i*_ denote the cell population at time, *t* = 0, and at any time, *t* > 0, respectively, with *i* = 1, …, *n*. Let *i* = 1 denote the compartment for the G2-phase cells, then (25) and (34) become

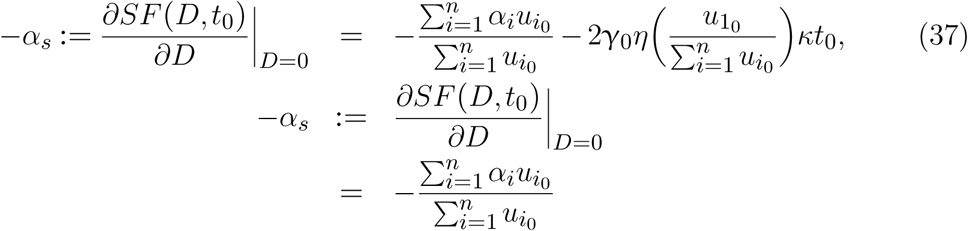

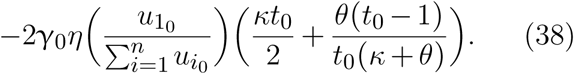

This implies that an increase in the number of model compartments only affects direct radiation effects.

In the next section, we will validate these derivations against the data for the *α*_*s*_ of the ten different asynchronous cell lines and three different synchronous cell lines used earlier in Section 3. We are interested in computing the initial slope of the SF curve using formula (26) and comparing it with the data in Joiner et al. [11].

## 5. Validation of the analytical derivations of the 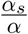 index

In this section, we validate the analytical results of the previous section against SF data of both synchronous and asynchronous cell lines. In particular, we estimate the initial slope of the SF curve using formula (26) and compare it to the data in Figure 2 of [11], which is the plot of 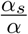 vs. the surviving fraction at 2 Gy *SF*_2_. The values of *α*_*s*_ computed using formula (26) with parameter values in Table 1 and 2 are recorded in Table 3 for asynchronous cells and Table 4 for synchronous cells, respectively. The third row contains the value of the 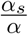 index for each cell line and cell cycle phase, as the case may be. The *α* values used in the computation of 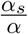 index are from Tables 1 and 2 while *SF*_2_ values are taken from respective literature.

**Table 3:** Computed values of 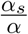 and literature values of the *SF2* for asynchronous cell lines. The mean ± SD of *α*_*s*_ are computed from (26) using the estimated parameters in Table 1; values of 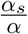 are computed using the mean of *α*_*s*_ and the values of *α* from Table 1; *SF*_2_ data are taken from the respective literature.

|  | MR4 | PC3 | V79 | V79<br>ox | V79<br>hyp | A549 | HT29 | U1 | T98G | RWPE1 |
| --- | --- | --- | --- | --- | --- | --- | --- | --- | --- | --- |
| $\alpha_s$ | $2.727 \pm 0.774$ | $0.703 \pm 0.448$ | $0.470 \pm 0.280$ | $0.394 \pm 0.3032$ | $0.289 \pm 0.215$ | $0.484 \pm 0.362$ | $0.222 \pm 0.162$ | $0.353 \pm 0.186$ | $1.858 \pm 1.057$ | $3.230 \pm 0.810$ |
| $\frac{\alpha_s}{\alpha}$ | 64.94 | 6.241 | 2.981 | 3.294 | 3.368 | 4.068 | 6.784 | 28.24 | 13.78 | 37.15 |
| $SF_2$ | 0.483 | 0.643 | 0.674 | 0.667 | 0.847 | 0.682 | 0.775 | 0.910 | 0.748 | 0.607 |

**Table 4:** Computed values of 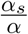 and literature values of the *SF*_2_ for synchronous cell lines. The mean ± SD of *α*_*s*_ are computed from (26) using the estimated parameters in Table 2; values of 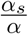 are computed using the mean of *α*_*s*_ and the values of *α* from Table 2; *SF*_2_ data are taken from the respective literature.

|  | V79<br>G1 | V79 S | V79<br>G2 | T98G<br>G1 | T98G<br>S | T98G<br>G2 | U373<br>G1 | U373 S | U373<br>G2 |
| --- | --- | --- | --- | --- | --- | --- | --- | --- | --- |
| $\alpha_s$ | $0.253 \pm 0.016$ | $0.155 \pm 0.067$ | $2.779 \pm 2.017$ | $1.157 \pm 0.549$ | $1.335 \pm 0.491$ | $5.890 \pm 4.202$ | $0.028 \pm 0.017$ | $0.120 \pm 0.054$ | $5.203 \pm 3.195$ |
| $\frac{\alpha_s}{\alpha}$ | 1.012 | 1.824 | 9.172 | 8.900 | 10.26 | 28.05 | 1.400 | 1.200 | 32.52 |
| $SF_2$ | 0.63 | 0.77 | 0.53 | 0.83 | 0.67 | 0.61 | 0.61 | 0.70 | 0.49 |

The numerical validation of formula (26) against data is shown in Figure 7. This Figure shows the relationship between the 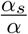 index and the surviving fraction at 2 Gy, *SF*_2_. In this figure, we superimpose two sets of data. First, we plot the model results from Tables 3 and 4, and then superimpose experimental results from [11]. The plot in [11] contains 33 different cell lines. Although Figure 7 has fewer cell lines, we observe a good qualitative agreement between model results and a subset of the data from [11]. This affirms that the values of *α*_*s*_ computed from formula (26) agree with the existing data.

**Figure 7:**
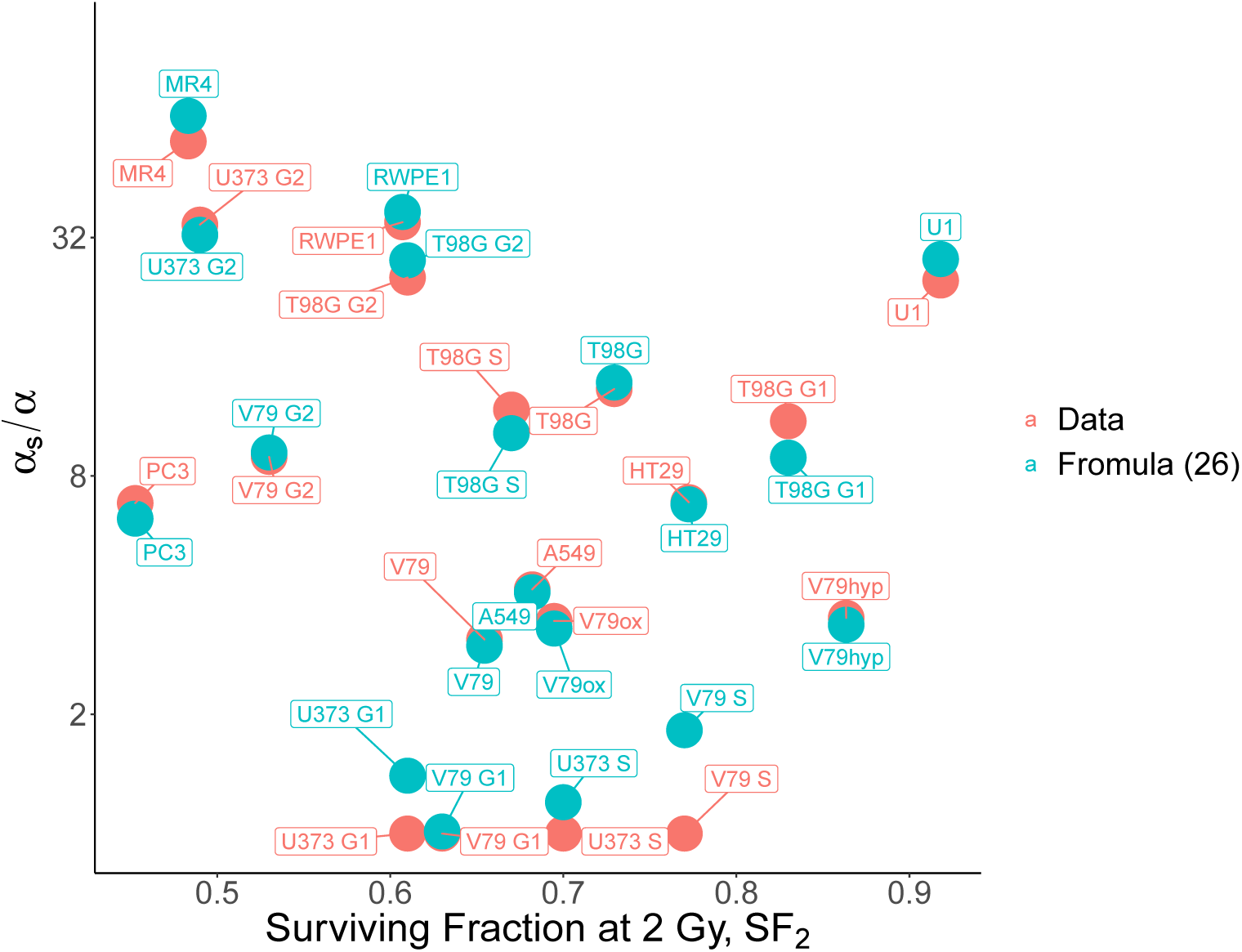
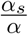 index measured at low doses of radiation against the surviving fraction measured at 2 Gy, *SF*_2_. Red bullets represent data of different synchronous and asynchronous cells line from [11, 19, 30]; green bullets represent the computational results from formula (26).

## 6. Relationship between the 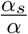 index and radioresistance at 2 Gy

As noted earlier, considerable attempt has been made in the past to describe the relationship between the degree of HRS exhibited in a cell and its radioresistance at 2 Gy. In [14], Lambin et. al found that cells that exhibit a higher degree of HRS also demonstrate a significant increase in radioresistance at 2 Gy. It was later found in [11] that more cell lines data do not agree with this positive relationship.

In this section, we will analytically derive this relationship and also numerically explore the relationship for better understanding of the radioresistance at 2 Gy vs. HRS exhibited in a cell. Fortunately, the derivation is simplified because of the earlier work in Section 4.

From the LQ model, we derive;

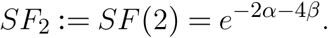

This implies that

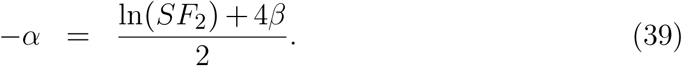

From (26), we have

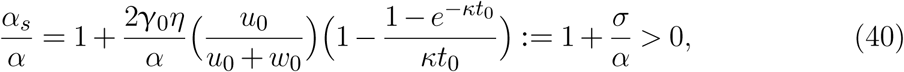

where

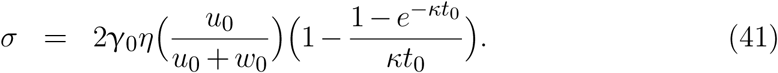

Henceforth, the parameter *σ* in (41) will be referred to as the *intrinsic radiosensitivity* of a cell. This terminology makes sense since it is the indirect radiation effect that determines the degree of HRS a cell exhibits. Combining (39) and (40) gives

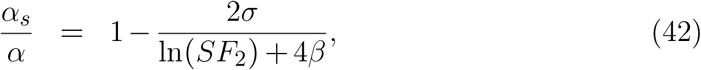

which depends on *σ*, the cell’s intrinsic radiosensitivity, *β*, the rate at which binary misrepair of pairs of DSB from different radiation tracks lead to lethal lesions, and *SF*_2_, the surviving fraction at 2 Gy. Our interest is to understand how these three parameters affect the data in Figure 7 using (42).

As noted earlier that Lambin et. al [14] suggested that a universal relationship between the 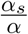 index and *SF*_2_ should exist; and that the hypothesis was not supported by the study of more cell lines. Here we find that such a relationship does exist, however, it depends on two cell-specific parameters, the intrinsic radiosensitivity *σ* and the direct radiation sensitivity *β*. In Figure 8, we show that the cell lines we examined do fall onto the curve defined by (42). Since it is hard to visualize a plot in 4-dimensions, we fix either *σ* or *β* in order to contour plot function (42). It is worth noting that function (42) has a vertical asymptote at *SF*_2_ = *e*^−4*β*^ for each value of *β*. Figure 8a is a surface plot of (42) for *σ* = 2. This plot is an increasing manifold in both *β* and *SF*_2_, respectively. The white dots represent different cell lines lying on the manifold based on their corresponding values for *β* and *SF*_2_. Similarly, the surface plot in Figure 8b for a fixed *β* = 0.01 *Gy*^−2^ also shows increasing trends both in *σ* and *SF*_2_ and a good match with the white dots corresponding to different cell lines. Note that if we project these plots onto the (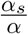, *SF*_2_) plane, we will get a 2-dimensional plot similar to Figure 7, but in which the trends no longer are obvious. Hence, the trend suggested by Lambin et. al in [14] indeed holds across cell lines. In other word, if the data in Figure 7 was plotted in 3-dimension with *β* or *σ* as the third axis, we will have different cell lines lying on different layers of these manifolds depending on and increasing in both *β* and *σ*.

**Figure 8:**
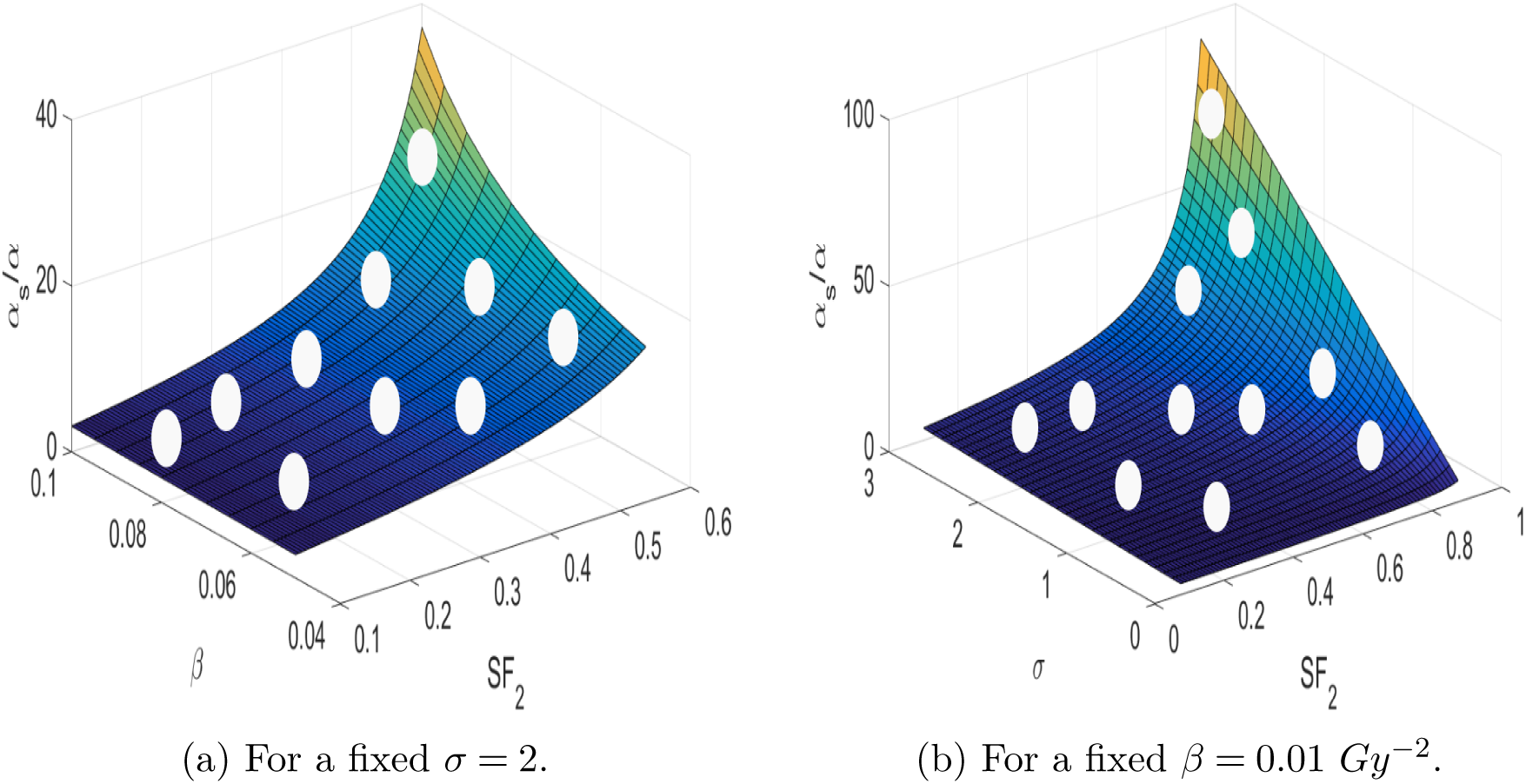
Surface plot of the relationship (42) between the 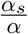 index and *SF*_2_, the surviving fraction at 2 Gy, at different values of *σ* and *β*, respectively. The white dots represent different cell lines depending on their *σ* and *β* values. (a) The surface plot for a fixed *σ* = 2; (b)The surface plot for a fixed *β* = 0.01 *Gy*^−2^.

## 7. Discussion

The phenomenon of low-dose hyper-radiosensitivity has been observed across many different kind of cell lines. Several malignant cancer cells like glioma cells (T98G) and prostate cancer cells (PC3) are among the cells that exhibit this low-dose phenomenon. Many researchers [2, 5, 28, 31] are currently investigating how this phenomenon can be exploited to improve the effectiveness of cancer radiotherapies. The major obstacle to this idea is the superficiality in the level of understanding of this low-dose phenomenon.

It is worth noting that there are other mechanisms, which are not cell cycle-based, that can trigger the low-dose HRS phenomenon. For example, the mechanism of bystander effects have been implicated in the HRS phenomenon in [26, 27, 22, 25]. Moreover, Mothersill et al. [23] have shown that both the mechanisms of bystander effects and G2 checkpoint are two mutually exclusive cellular events.

The most common approach for measuring this low-dose phenomenon in a cell is by comparing the initial slope of its SF curve *α*_*s*_, to that of its initial slope of the SF curve *α*, described by the LQ model. That is, by determining the 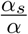 index. We understand that this index quantifies the degree of HRS in a cell and must relate to some of the molecular cascades or signaling pathways implicated in the occurrence of HRS phenomenon [12, 13, 4]. However, the specific form of this relationship is yet unknown.

In this paper, we attempt to unravel this relationship by building a system of ODE that captures the interaction between radiation exposure and cells in different cycle phases. Our model is able to fully capture the observed HRS phenomenon in ten different asynchronous cell lines and three different synchronous cells in different cell cycle phases.

The derivation in Section 4 of *α*_*s*_, the initial slope of the SF curve, reveals the dependence of *α*_*s*_ on the proportion of cells in the G2 phase, the rate of mitotic catastrophe experienced by damaged G2 cells, the rate of radiation-induced cell damage, the rate of damage repair, and the duration of radiation exposure. This is consistent with experimental observations that damaged G2 cells which evade the early G2 checkpoint to proceed to mitosis undergo mitotic catastrophe, which results in the increased sensitivity of some cells at low doses. Although, our result is not surprising, the analytic quantification of this dependence is novel and we observe a good qualitative agreement between this dependence and experimental data as shown in Figure 7.

In the case where the duration of radiation exposure is long enough to accommodate DNA repair. We observe that the repair rate *θ*, and the rate of mitotic catastrophe *κ*, contribute differently to the degree at which a cell exhibits HRS. Our model shows that for small mitotic catastrophe rate *κ*, the slope *α*_*s*_ of the SF curve is more sensitive to changes in the DNA repair rate *θ*, than for larger values of *κ*.

In these analytic relationships, we realize that there is no dependence on any of the parameters in checkpoint function, *c*(*D*). This further supports the evidence that the low-dose HRS is a result of ineffective checkpoint activity while the IRR is a consequence of checkpoint activation. The model developed in this paper can also be used to study the effect of checkpoint activity on the low dose IRR. However, this will be outside the scope of consideration since we are only exploring the phenomenon of HRS in this paper.

Another important concern in (26) is that it does not consider the effect of cells that were damaged in non-G2 phases but have progressed into G2 phase with unrepaired damage. For instance, suppose only cells in non-G2 phases are irradiated. In a matter of time, there will be more cells entering into the G2 phase, some of which will carry over damage from other cycle phases. It is then reasonable to ask if these damaged cells in G2 phase also contribute to the mechanism underlying the phenomenon of HRS. In response to this concern, there are biological experiments [12, 13] that have shown that the checkpoint mechanisms that control the mitosis of G2 cells that carry over damage from other phases is different from the mechanism that controls the cells that sustain radiation damage right in the G2 phase. The checkpoint that controls the former is called the Sinclair Checkpoint while the checkpoint that controls the later is called the Early G2 Checkpoint. An experiment by Krueger et. al. in [12] has shown that the Sinclair checkpoint does not contribute to the HRS phenomenon. This is also confirmed in the formula for the initial slope of the SF curve in (27).

In order to investigate the relationship between 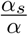 index and the surviving fraction at 2 Gy *SF*_2_, we derive an analytical formula which is a function of three parameters, namely; *SF*_2_, the surviving fraction at 2 Gy, *σ*, the intrinsic radiosensitivity, and *β*, the rate of binary misrepair of pairs of DSB forming lethal lesions. We found that for each value of either *σ* or *β*, the increasing trend in *SF*_2_ is preserved by a manifold increasing in either both *σ* and *SF*_2_ or *β* and *SF*_2_. Thus, the cells with higher 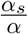 index can also have higher *SF*_2_ value depending on the corresponding values of *σ* and *β*. This result generalizes Lambin’s result in [14] to any cell lines.

The mathematical derivations in this paper do not account for the situation when the SF of cells is not measured immediately after radiation exposure. We consider only the cases when the SF is measured immediately after radiation because it simplifies the mathematical technique for the derivations. The actual experimental procedure consists the radiation of cells and a post-radiation incubation period within which the survived cells can be detected. These are separate experiments and our model can only describe the event until the end of radiation exposure. Our model then assumes that the outcome of the second experiment involving incubation can be approximated by computing the SF from our model immediately after an exposure. This assumption was justified by the qualitative agreement between our model computational results and the experimental data.

The current understanding of the low-dose HRS is still limited in the context of cell lines. There is a need to understand the connection to low-dose radiation in the context of human tissue. Such an understanding would have an immense implication on the current radiotherapy practices and perhaps could point the way to treatment strategies not yet considered.

## Acknowledgements

The authors are grateful for the feedback from the Journal Club of the Collaborative Mathematical Biology Group at the University of Alberta. OO is supported by the University of Alberta President’s International Doctoral Award and Queen Elizabeth II Graduate Scholarship - Doctoral level, GdeV and TH are supported by the Natural Sciences and Engineering Research Council of Canada (NSERC).

